# Nanoscale PI(4,5)P_2_ partitioning driven by FGF2 oligomerization triggers membrane pore formation

**DOI:** 10.1101/2025.09.11.675533

**Authors:** Alessandra Griffo, Daniel Beyer, Jaime Fernández Sobaberas, Melanie König, Zeynep Kilcan, Manpreet Kaur, Eleonora Margheritis, Katia Cosentino, Petra Riegerová, Martin Hof, Radek Šachl, Walter Nickel, Fabio Lolicato

**Author notes:** These authors contributed equally to this work (Alessandra Griffo, Daniel Beyer, Jaime Fernández Sobaberas).

## Abstract

Fibroblast Growth Factor 2 (FGF2) is a potent tumor cell survival factor. Unlike most secreted proteins, FGF2 lacks a signal peptide and is exported by an unconventional secretory pathway, bypassing the ER/Golgi system. A key event of this unusual secretory mechanism is FGF2 recruitment at the inner plasma membrane leaflet mediated by the phosphoinositide PI(4,5)P₂. This interaction induces FGF2 oligomerization which, in turn triggers the formation of a transient lipidic membrane pore with a toroidal structure. FGF2 oligomers populating such membrane pores are captured and disassembled at the outer plasma membrane leaflet through interactions with heparan sulfate chains of the cell surface proteoglycan GPC1, resulting in full translocation of FGF2 across the plasma membrane. Once FGF2 appears on cell surfaces, it engages in signaling complexes initiating autocrine or paracrine signaling. Here, using a multidisciplinary and multiscale approach, we show that FGF2 self-assembles into oligomers that induce lateral PI(4,5)P₂ partitioning and membrane remodeling. This process depends on PI(4,5)P₂ reaching a critical local threshold concentration that, due to its non-bilayer properties destabilizes the bilayer, facilitating pore formation and FGF2 translocation across the plasma membrane. Our findings highlight a fundamental principle of how protein oligomerization can induce lateral partitioning of membrane lipids that in turn leads to a drastic membrane remodeling event: the transformation of the lipid bilayer into a toroidal membrane pore. The underlying mechanism is likely to be applicable to a wide range of membrane lipid interacting proteins that induce membrane remodeling in other cellular processes.

## Introduction

Fibroblast Growth Factor 2 (FGF2) is a potent mitogen with prominent roles in the development of the vascular system and wound healing (1,2). In addition, under pathophysiological conditions, FGF2 serves as both a mediator of tumor-induced angiogenesis and acts as a survival factor for cancer cells inducing chemoresistance in the treatment of for example acute myeloid leukemia. The latter activity is mediated by FGF2’s ability to block apoptosis through an autocrine secretion-signaling loop (3,4).

Unlike the majority of secretory proteins containing N-terminal signal peptides for ER/Golgi-dependent protein secretion, FGF2 is transported into the extracellular space by direct translocation across the plasma membrane (5–7). The entry point of this unconventional secretory pathway of protein secretion is defined by the initial recruitment of FGF2 at the inner plasma membrane leaflet mediated by the α1 subunit of the Na,K-ATPase, contacting FGF2 in a region on its protein surface containing K54, K60 and C77 (8,9). A subsequent interaction involves Tec kinase, resulting in tyrosine phosphorylation of FGF2 at Y81 (10,11). Following initial accumulation of FGF2 at the inner leaflet mediated by the Na,K-ATPase and Tec kinase, FGF2 is handed over to the phosphoinositide PI(4,5)P_2_ whose head group (IP_2_) binds a highly specific binding pocket of FGF2 that contains the surface residues K127, R128 and K133 (12–15). In addition, multiple low affinity interactions of FGF2 with PI(4,5)P_2_ molecules have been observed in molecular dynamics simulations, consistent with the overall basic isoelectric point of 9.6 of FGF2 (15). The interaction with PI(4,5)P_2_ triggers FGF2 to oligomerize (12,13) with disulfide-bridged FGF2 dimers serving as building blocks for higher FGF2 oligomers (16). Disulfide bridges in FGF2 oligomers are homogenously formed from C95 residues that were shown to be essential for both FGF2 oligomerization in vitro and FGF2 secretion from cells into the extracellular space (14,16). The function of Tec kinase-mediated tyrosine phosphorylation of FGF2 that occurs on Y81 in close proximity to the C95-C95 disulfide bridge in FGF2 dimers may facilitate oxidative FGF2 dimerization and/or protect such dimers against reduction of the disulfide bridge and therefore conversion back into monomers at the cytoplasmic leaflet of the plasma membrane. This may explain the role of Tec kinase in facilitating the efficient secretion of FGF2 (10,11).

In previous studies, PI(4,5)P_2_-dependent FGF2 oligomerization has been identified to trigger the formation of a lipidic membrane pore with a toroidal structure (6,13–16), a process that has recently been demonstrated to be facilitated by the asymmetric transbilayer distribution of PI(4,5)P_2_ (17). Upon opening, FGF2 oligomers were proposed to populate the inner hydrophilic space of the produced membrane pore, thereby getting exposed at the outer leaflet of the plasma membrane. Full FGF2 membrane translocation was shown to be completed by a further essential component of this unusual mechanism of protein secretion, Glypican-1 (GPC1) (5,18,19). GPC1 is a proteoglycan localized on cell surfaces whose membrane proximal heparan sulfate chains contain high affinity binding sites for FGF2 (18,19). Once PI(4,5)P_2_-bound FGF2 oligomers localized within the lipidic membrane pore become accessible at the outer plasma membrane leaflet, GPC1 captures and disassembles FGF2 oligomers, most likely into disulfide-bridged dimers (16), outcompeting PI(4,5)P_2_ based on overlapping binding sites in FGF2 with K133 being essential for FGF2 interactions with both PI(4,5)P_2_ and heparan sulfate chains (15). The nature of FGF2 interactions with PI(4,5)P_2_ (K_D_ ∼2 µM; (12,15,20)) versus heparan sulfates (K_D_ ∼50 nM; (21–23)) was found to be mutually exclusive, an observation that has been proposed to represent the molecular basis for vectorial translocation of FGF2 from the cytoplasmic side across the plasma membrane into the extracellular space (5).

In terms of the lateral organization of the FGF2 membrane translocation machinery, we have hypothesized the existence of nanodomains that house all of its components in ordered membrane domains (24,25). In support of this, GPC1 is a proteoglycan that is embedded into the plasma membrane via a GPI anchor and accordingly found exclusively in ordered domains (5,19). In addition, we found cholesterol to be enriched in the vicinity of interactions between FGF2 and PI(4,5)P_2_ along with observations of high cholesterol levels supporting both increased FGF2 binding to PI(4,5)P_2_ and increased FGF2 secretion efficiency from cells (26). A pre-assembled organization of the FGF2 membrane translocation machinery in specialized nanodomains may also be one of the underlying principles of the extremely fast FGF2 membrane translocation kinetics with time intervals of about 200 ms for a single membrane translocation event as measured in intact cells by single-molecule real-time TIRF microscopy (27). Following membrane translocation, FGF2 dimers remain bound to heparan sulfate chains on cell surfaces, either being stored or engaging in the formation of ternary signaling complexes activating FGF receptors for either autocrine or paracrine signaling (1,2).

In the current study, based on in vitro approaches, imaging experiments in living cells and multiscale molecular dynamics simulations, we reveal that the functional oligomeric state of FGF2 required to form membrane pores is a dynamic one, represented by tetramers and hexamers. Due to multiple PI(4,5)P_2_ molecules interacting with a single FGF2 molecule, FGF2 oligomerization into rings of tetramers and hexamers leads to a substantial local accumulation of PI(4,5)P_2_ molecules in a highly spatially constrained membrane surface area at the inner plasma membrane leaflet. Since PI(4,5)P_2_ is a non-bilayer lipid with an asymmetric transbilayer distribution (28), we propose that its local clustering into PI(4,5)P₂-enriched domains promotes membrane remodeling, driving the transition from a stable bilayer to a toroidal lipid pore. This study not only reveals the mechanistic basis of how FGF2 can physically traverse the plasma membrane as part of its unconventional pathway of secretion, but it furthermore has also fundamental mechanistic implications as a general principle for the potential of lipid interacting proteins to laterally sort membrane lipids through protein oligomerization, initiating membrane remodeling events in a wide range of physiological processes.

## Results

### Quantification of the oligomeric state of FGF2 on the membrane surface of GUVs

To investigate fibroblast growth factor 2 (FGF2) oligomerization in giant unilamellar vesicles, we reconstituted vesicles with a composition designed to approximate the major lipid constituents of the plasma membrane. The lipid mixture contained seven components in defined molar ratios, supplemented with biotinylated phosphatidylethanolamine for surface immobilization (the detailed lipid composition is provided in **Table 1** and described in the Materials and Methods section). The vesicles were incubated with fluorescently labeled FGF2-Y81pCMF-eGFP (13,15), and FGF2 oligomerization was quantified using dual(+1) fluorescence correlation spectroscopy (FCS). This method was originally developed to probe FGF2 membrane insertion and oligomerization (29,30). In contrast to conventional FCS, the dual(+1) approach allows selective analysis of vesicles that contain functionally oligomerized, membrane-inserted FGF2 complexes. This is possible because insertion of functional FGF2 into the membrane consistently leads to vesicle permeabilization (15,17). Permeabilized vesicles can thus be identified and isolated for further brightness analysis, ensuring that only functionally relevant oligomers are studied. This filtering step was critical, as it excludes vesicles dominated by non-specific protein aggregation, which would otherwise bias the analysis toward artificially high oligomer sizes.

**Table 1.** Lipid compositions of GUVs used for dual(+1)-FCS measurements.

| Abbreviation | Full name | mol% |
| --- | --- | --- |
| PC | Phosphatidylcholine | 33 |
| PE | Phosphatidylethanolamine | 9 |
| PS | Phosphatidylserine | 5 |
| PI | Phosphatidylinositol | 5 |
| SM | Sphingomyelin | 15 |
| PIP(4,5)2 | Phosphatidylinositol-4,5-bisphosphate | 2 |
| CHOL | Cholesterol | 30 |
| Biotinyl-PE* | Biotinylated phosphatidylethanolamine | 1 |
| DGS-NTA | 1,2-dioleoyl-sn-glycero-3-[(N-(5-amino-1-carboxypentyl)iminodiacetic acid)succinyl]<br>(nickel salt) | 2 |

\* GUVs contained either 2 mol% PI(4,5)P<sub>2</sub> or 2 mol% DGS-NTA(Ni).

A total of 67 giant unilamellar vesicles were analyzed under these conditions. The distribution of the oligomeric states of FGF2 is shown in **Figure 1**, with the histogram plotted alongside a violin plot of protein surface concentration. The oligomer size distribution displayed a pronounced peak to four to six subunits, accompanied by a long tail extending toward higher aggregation numbers. Across the population, the median oligomer size was 5.1, with a mean of 5.7 ± 3.12. Quantification of membrane association revealed that FGF2 bound to the GUV membrane surfaces at a median protein surface concentration of 0.81 nmol/m² (mean 0.83 ± 0.329 nmol/m²). This corresponds to an average protein-to-lipid ratio of 1:2830 under the tested conditions. These findings indicate that FGF2 associates with membranes at relatively low surface densities but, once bound, preferentially assembles into tetrameric and hexameric complexes. The overall distribution thus reflects a dominant tendency toward mid-sized oligomers, accompanied by a smaller fraction of both dimers and higher stoichiometries across individual vesicles.

**Figure 1.**
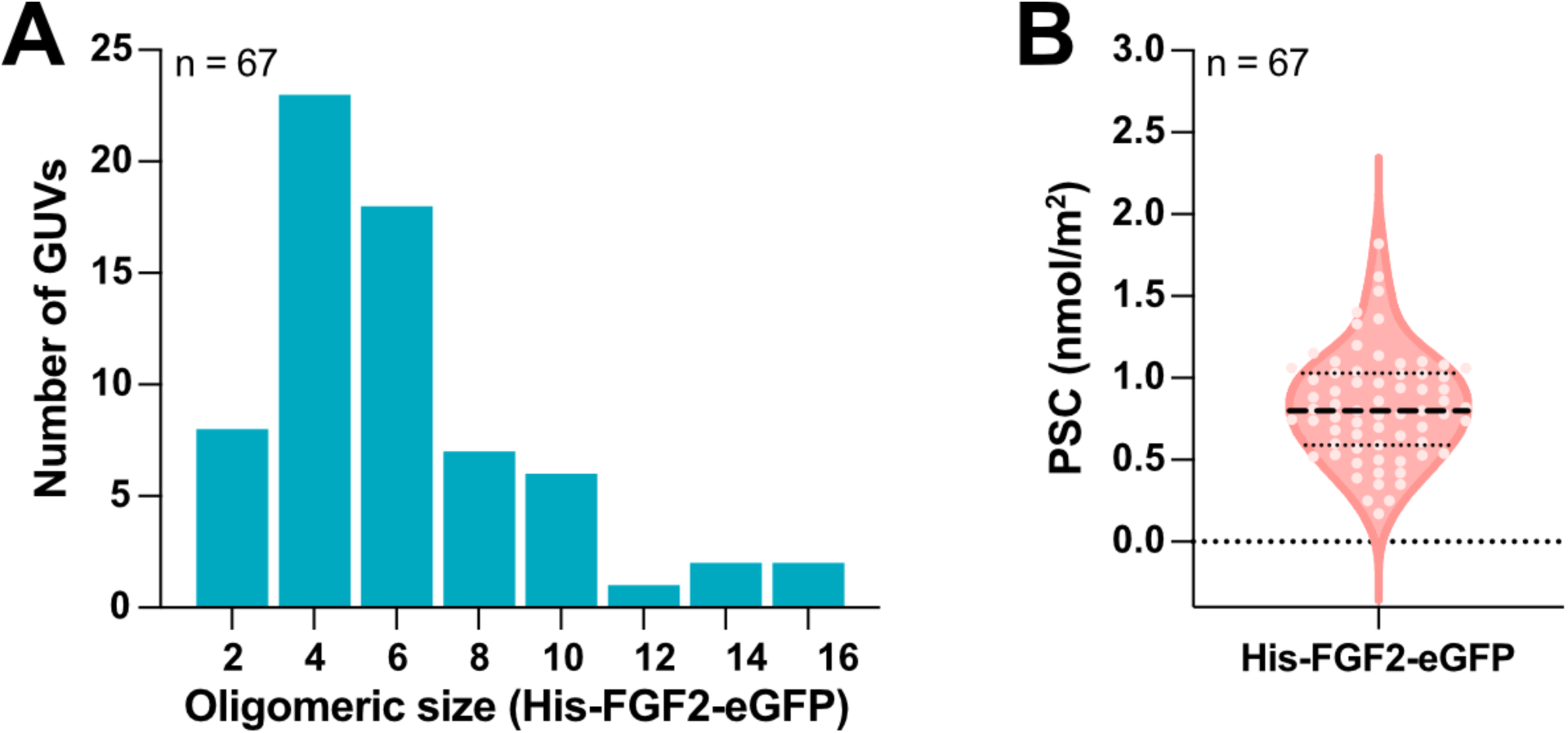
Brightness analysis determining the range of oligomeric species of FGF2 at the surface of GUVs employing dual(+1) fluorescence correlation spectroscopy. His-FGF2-eGFP oligomer size distribution (a) and the corresponding distribution of membrane surface concentration of His-FGF2-eGFP (b) measured on 67 permeabilized GUVs by dual(+1)-FCS.

### Quantification of the oligomeric state of FGF2 at the inner plasma membrane leaflet of living cells

As documented in **Figure 1** in the previous section, FGF2 tetramers and hexamers could be identified to represent the functional oligomeric species of FGF2 in biochemical reconstitution experiments by analyzing membrane pore formation in GUVs containing PI(4,5)P_2_. In the cell-based experiments shown in **Figure 2**, we aimed at quantifying the distribution of oligomeric species of FGF2 at the inner plasma membrane leaflet of intact cells. Employing total internal reflection fluorescence (TIRF) microscopy with single-molecule resolution (27,31,32), we conducted a brightness analysis of FGF2-GFP oligomers to determine the subunit stoichiometry of FGF2 oligomers in the vicinity of the inner plasma membrane leaflet in living CHO cells. As a control, we compared the wild-type form of FGF2-GFP with FGF2-C95A-GFP, a variant form of FGF2 that has been demonstrated to fail in forming functional oligomers, to be deficient in membrane pore formation and to be incapable of being secreted from cells (16). The analyses were based on stable CHO cell lines in which expression of the FGF2-GFP fusion proteins indicated can be induced with doxycycline (16,26,33). Cells were seeded on microscopy chips and FGF2-GFP expression was induced at low levels to allow for single-molecule detection. Using TIRF microscopy, FGF2-GFP oligomers could be visualized as bright fluorescent spots associated with the plasma membrane (**Figure 2 A-B**). In those images, it was immediately apparent that FGF2-C95A-GFP spots appeared less bright than FGF2-GFP spots, indicating that the C95A variant does not assemble into higher order oligomers. A detailed stoichiometry brightness analysis revealed FGF2-GFP particles to represent a dynamic range of higher order oligomers (**Figure 2 C**). By contrast, the FGF2-C95A-GFP variant form showed a narrow distribution of oligomeric species that was consistent with the lower brightness of the particles observed to be associated with the plasma membrane (**Figure 2 D**). Using the average brightness intensity of a monomeric GFP variant targeted to the plasma membrane (TMD-mEGFP) used as a reference molecule for calibration based on the brightness of one EGFP unit, we estimated the subunit stoichiometry of both FGF2-GFP and FGF2-C95A-GFP oligomers at the plasma membrane of CHO cells. This analysis suggested that FGF2 dimers serve as building blocks for higher order FGF2 oligomers, the latter found to be represented mainly by tetramers and hexamers (**Figure 2 E**). FGF2-GFP monomers were undetectable. By contrast, tetramers and hexamers were nearly undetectable for FGF2-C95A-GFP. Instead, we found the majority of FGF2-C95A-GFP to be assembled into dimers and, to a lesser extent, existed as monomers. As previously reported for in vitro experiments (16), the data on FGF2-C95A-GFP confirm that the formation of a C95-C95 disulfide bridge is not an essential requirement for FGF2 dimer formation. However, the previously identified electrostatic interface, which mediates the transient formation of FGF2 dimers brings into close contact the thiol-containing side chains of two C95 residues into close contact, promoting the formation of covalent FGF2 dimers with the monomers linked via a disulfide bridge ^4^. The findings shown in **Figure 2 E-F** demonstrate that the formation of covalent C95-C95 bridged FGF2 dimers is essential for the formation of higher order FGF2 oligomers that form membrane pores and therefore represent the intermediates required for unconventional secretion of FGF2. The identification of tetramers and hexamers for FGF2-GFP is consistent with the results from biochemical reconstitution experiments shown in **Figure 1**, identifying these oligomeric species to be the functional units of FGF2 membrane translocation.

**Figure 2:**
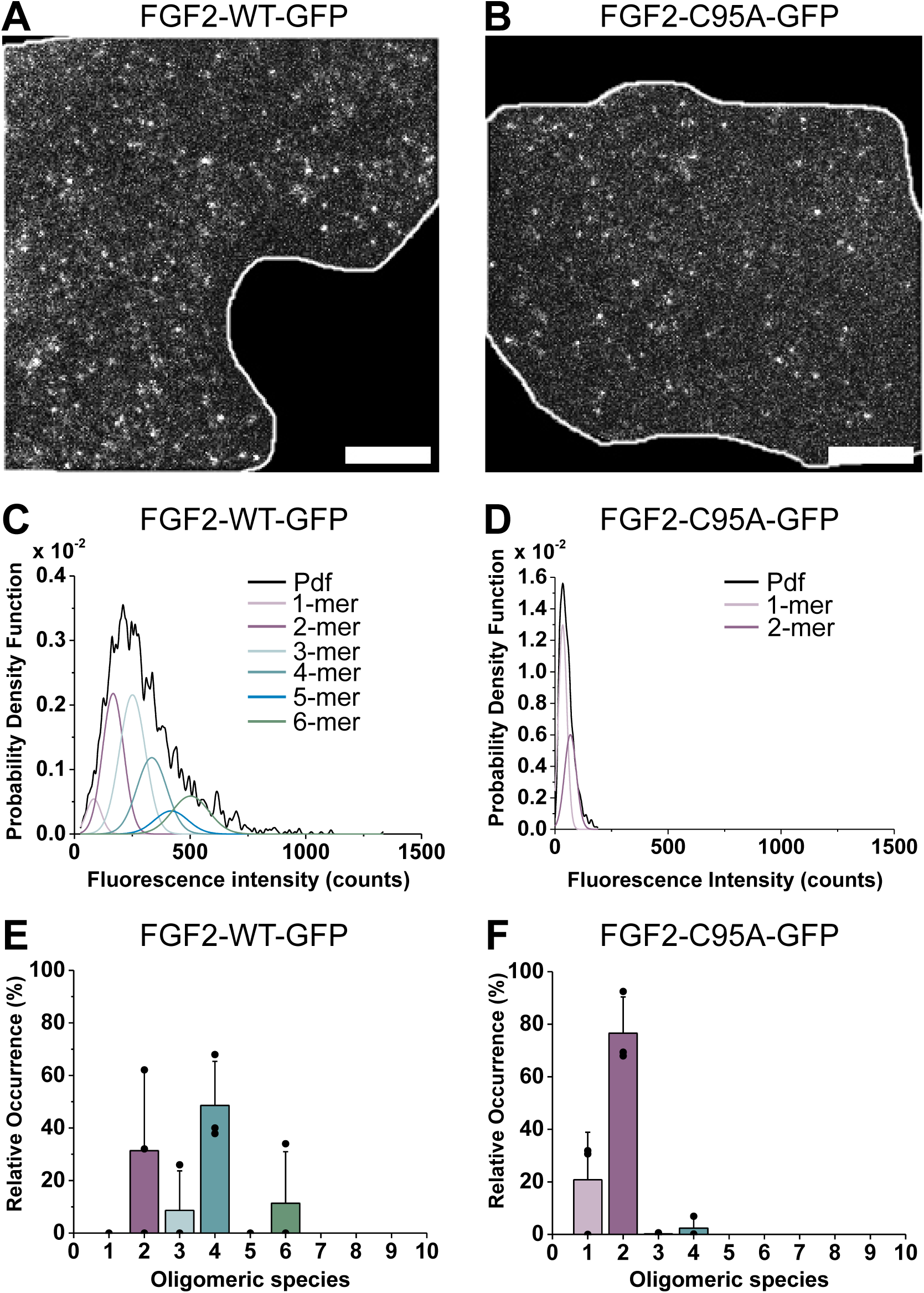
Brightness analysis determining the range of oligomeric species of FGF2 at the inner plasma membrane leaflet of intact cells employing TIRF microscopy with single-molecule resolution. A, B) Representative TIRF images of FGF2-WT-GFP (A) and FGF2-C95A-GFP (B) oligomers at the plasma membrane of CHO cells. Scale bar = 5 µm. C, D) Representative examples of fluorescence intensity distributions of FGF2-GFP oligomers for FGF2-WT-GFP (C) and FGF2-C95A-GFP (D). TIRF images as shown in panels A and B were subjected to a single-molecule brightness analysis using the Stoichiometry Analysis Software (31) (see Materials and Methods for details). Bright spots corresponding to FGF2-GFP oligomers were automatically detected. Brightness values of each individual bright spot were obtained. Brightness distributions of all values were plotted as probability density functions shown in black. Probability density functions were then fitted with a mixture of Gaussians generated based on the fluorescence intensity of the monomeric reference molecule (TMD-mEGFP) in the calibration data set, representing the range of possible oligomeric states FGF2-WT-GFP and FGF2-C95A-GFP can assume. Different oligomeric species are shown in different colours. (E, F) Relative occurrence in % of oligomeric species of FGF2-WT-GFP and FGF2-C95A-GFP plotted with the colour code used in panels C and D calculated from the area of each fitted Gaussian with SAS. The plotted data correspond to the average value from three independent experiments, each with a total of 40-50 cells and a minimum of a total of 1500 particles being analysed per experiment. Individual data points per experiment are indicated as black dots. Data were corrected for partial fluorescence of GFP in particles as described in Materials and Methods. Error bars correspond to the standard deviation calculated from three independent experiments.

### Atomistic molecular dynamics simulations of PI(4,5)P₂-dependent self-assembly of FGF2 hexamers

To study the self-assembly of FGF2 into oligomers, we performed atomistic molecular dynamics simulations using the CHARMM36m (34) force field and GROMACS-2023 (35). The proteins were placed on the surface of a POPC bilayer containing 30 mol% cholesterol and 5 mol% 18:0/20:4 PI(4,5)P_2_, with PI(4,5)P*₂* asymmetrically distributed between the leaflets. The PI(4,5)P_2_ concentration was increased compared to the ∼2 mol% typically found in plasma membranes to ensure that some PI(4,5)P_2_ molecules remained available in the bulk membrane, rather than being completely sequestered by FGF2. This choice was motivated by our previous observations showing that a single FGF2 monomer can simultaneously interact with up to four PI(4,5)P_2_ molecules (15).

We focused on the disulfide-linked C95–C95 FGF2 dimer, which represents the basic building block of higher-order oligomers (16). Each simulation system contained three dimers, since tetramers and hexamers were found to be the most abundant oligomeric species in in vitro experiments employing fluorescence correlation spectroscopy (**Figure 1**) and TIRF experiments with intact cells (**Figure 2**). We constructed 27 independent systems in which the three dimers were placed in different relative orientations, with an initial separation of 1 nm to allow weak intermolecular interactions (van der Waals and Coulomb cutoffs were set to 1.2 nm; **Figure 3 A**). Each dimer was pre-bound to two PI(4,5)P_2_ molecules positioned in the experimentally validated binding pockets (K127, R128, K133). Simulations were run for at least 3.9 *μs* each, resulting in a cumulative sampling of 138.25 *μs* (see **Table 2** for details). Across these simulations, multiple oligomeric arrangements emerged (**Figure S1**). The most frequently observed structures were L- and V-shaped tetramers which lead to a U-shaped hexamer in two cases (**Figure S1** R01 and R18 systems).

**Figure 3.**
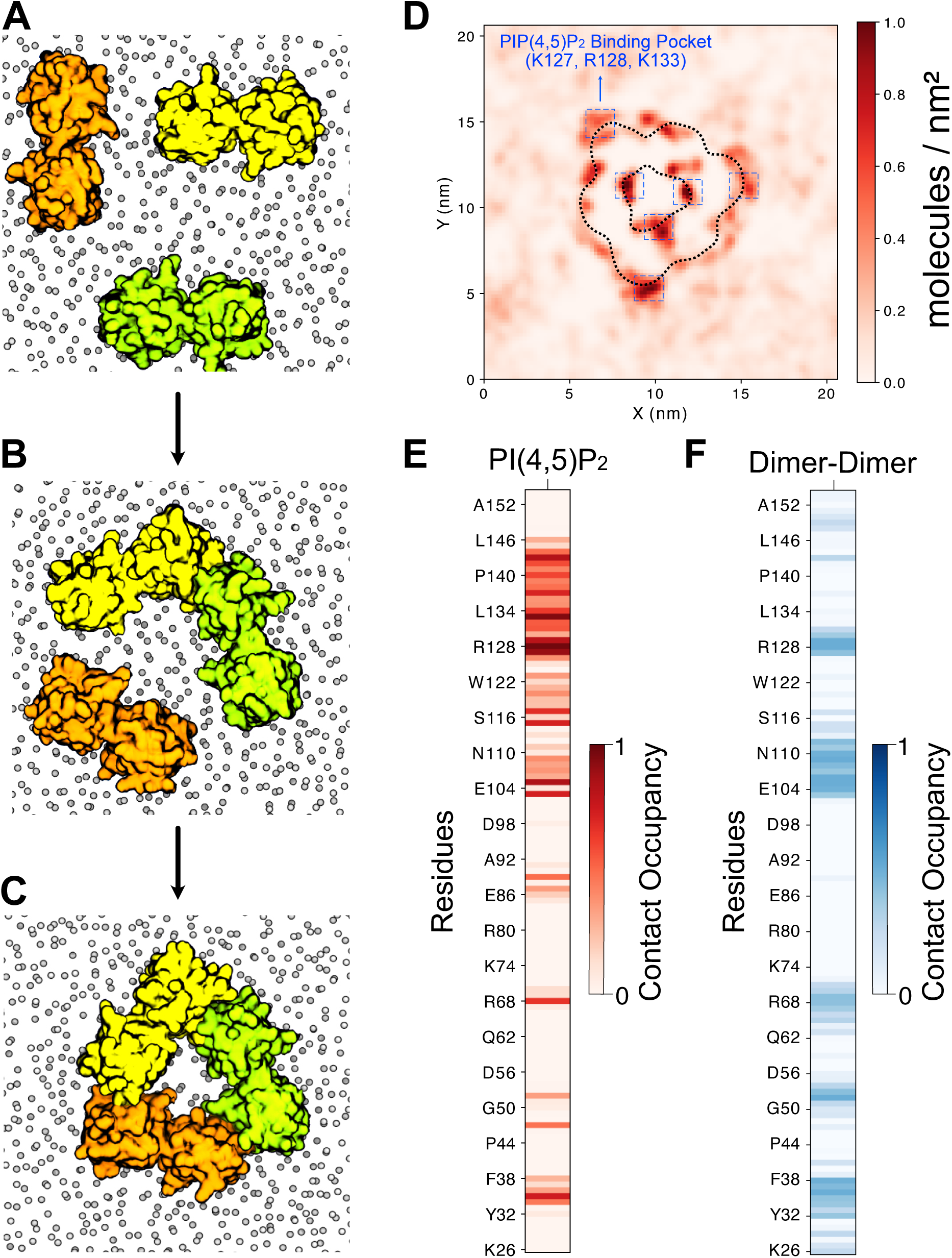
Atomistic molecular dynamics simulations of PI(4,5)P₂-mediated FGF2 hexamer self-assembly. (A) Representative initial configuration (one of 27 replicates) with three FGF2-C95–linked dimers embedded in a POPC:CHOL:PI(4,5)P₂ membrane. (B) Initial configuration showing the V-shaped tetramer, with a third dimer placed nearby at interacting distance. (C) Final configuration from a unbiased MD simulation yielding a ring-like hexameric assembly. (D) Two-dimensional PI(4,5)P₂ density map around the oligomer in a complex asymmetric membrane mimicking the plasma membrane. The map reveals PI(4,5)P₂ enrichment not only at the experimentally known binding pocket (highlighted in blue, residues K127, R128, K133) but also at additional sites. (E) Residue-wise contact occupancy with PI(4,5)P₂ lipids, showing preferential interactions. (F) Residue-wise contact occupancy at the dimer– dimer interface, identifying residues critical for oligomer stabilization. All analyses were performed on the final 1 μs of trajectory data and averaged across three independent simulations (n=3). A contact was defined as the distance between any atom of the two groups being ≤ 0.6 nm.

**Table 2.** Total simulation time of all coarse-grained self-assembly simulations with three dimers.

| System name | Time [μs] | System name | Time [μs] |
| --- | --- | --- | --- |
| R01 | 8,02 | R15 | 4,96 |
| R02 | 6,08 | R16 | 4,83 |
| R03 | 4,78 | R17 | 4,81 |
| R04 | 4,84 | R18 | 5,08 |
| R05 | 5,07 | R19 | 5,07 |
| R06 | 4,90 | R20 | 5,10 |
| R07 | 4,99 | R21 | 4,94 |
| R08 | 5,04 | R22 | 4,89 |
| R09 | 4,98 | R23 | 5,20 |
| R10 | 5,03 | R24 | 4,00 |
| R11 | 5,05 | R25 | 3,93 |
| R12 | 4,97 | R26 | 4,99 |
| R13 | 4,86 | R27 | 5,98 |
| R14 | 4,87 |  |  |

To test whether the closed hexameric state can be reached starting from a V-shaped tetramer, we placed a third dimer in close proximity at an interacting distance (**Figure 3 B**) and performed a 1.6-µs simulation. During the simulation, the third dimer initially interacted with one edge of the V-shaped dimer, forming a U-shaped hexamer. It then progressively engaged with the opposite extremity, ultimately completing the ring closure, as illustrated in **Figure 3C**. To further assess its stability, we embedded the hexamer in a more complex plasma membrane composition adapted from Schaefer and Hummer (36) (see **Table 3** for details), and performed three independent 5 μs long simulations (different initial velocities were used to assess independence). The hexamer remained intact throughout the total cumulative time of 15 μs. Importantly, the closed hexamer recruited PI(4,5)P_2_ not only at the canonical binding pockets but also at newly formed sites located at dimer–dimer interfaces (**Figure 3 D–F**). Such PI(4,5)P_2_-mediated stabilization was also observed for Gasdermin-D proteins, where ring-like oligomers were also stabilized by interface-associated PI(4,5)P_2_ lipids (36). Our contact analysis revealed both persistent PI(4,5)P_2_ binding and stable protein–protein interactions at the dimer–dimer interfaces (**Figures 3E–F**).

**Table 3.**
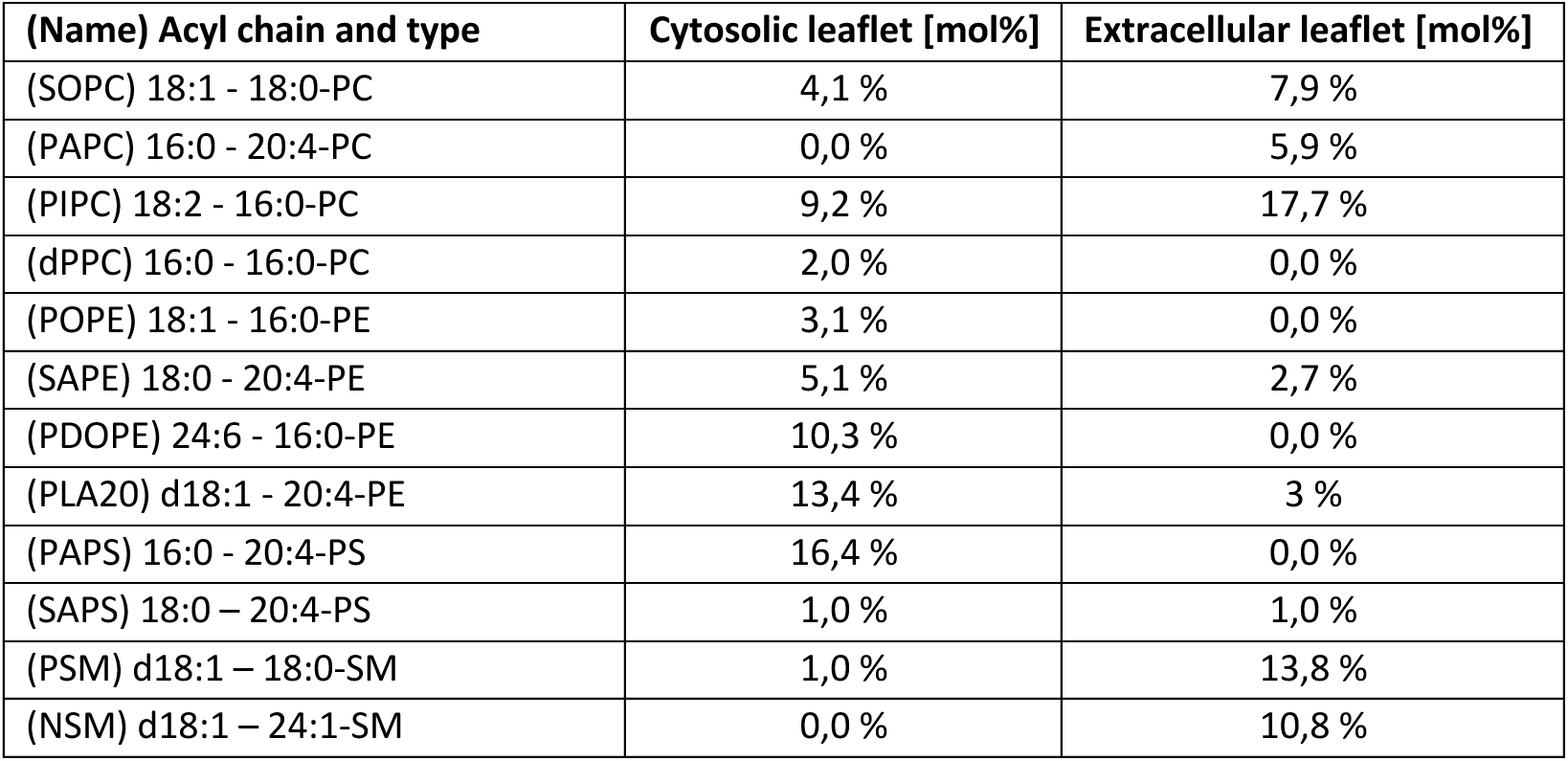
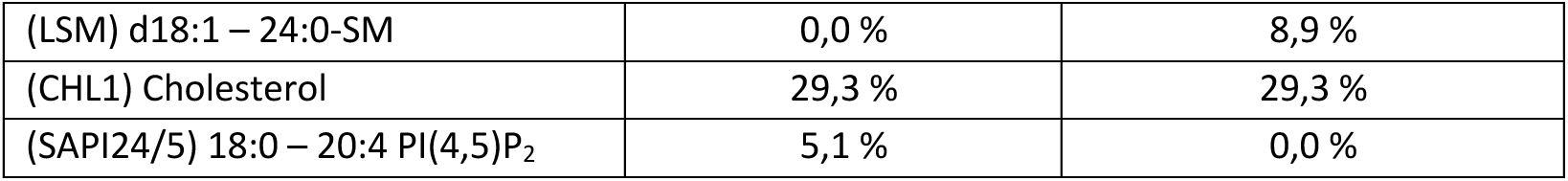
Initial lipid compositions of the asymmetric plasma membrane–like system (PM) adapted from Schaefer and Hummer(36).

Because lipid diffusion in atomistic simulations is intrinsically slow, and the accumulation of PI(4,5)P_2_ molecules around the hexamer further hinders sampling, we converted the atomistic hexamer model to the Martini 2.2 coarse-grained representation. This enabled us to efficiently probe lipid enrichment beneath the FGF2 hexamer at extended timescales.

**Figure S1.**
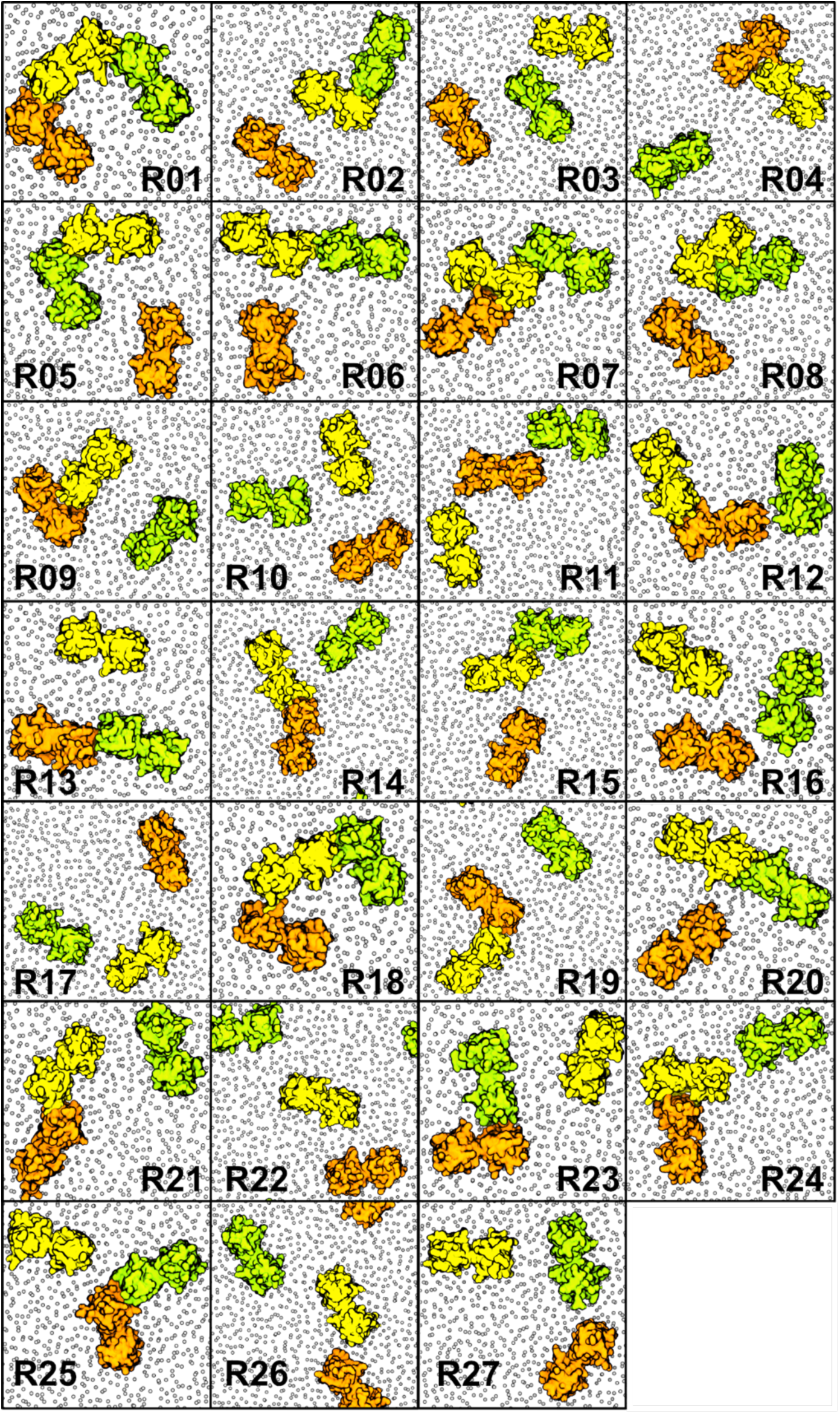
Final structural states observed in FGF2 self-assembly simulations. Shown are the final configurations from 27 independent molecular dynamics simulations of the self-assembly of three FGF2 dimers. Each simulation was initiated with a different relative orientation of three FGF2-C95–linked dimers and was run for at least 3.9 μs. Exact simulation lengths are provided in **Table 2** of the Materials and Methods section.

### FGF2 hexamers induce PI(4,5)P₂ clustering and membrane remodeling

To investigate the effect of the FGF2 hexamer on the plasma membrane, we converted the self-assembled ring-shaped FGF2 oligomer into a Martini 2 (37) coarse-grained model and positioned it on the cytoplasmic (inner) leaflet of an asymmetric plasma membrane containing 13 lipid species, distributed asymmetrically following the composition reported by Schaefer and Hummer (36) (**Figure 4 A** and **Table 3** for details). We specifically used Martini 2 rather than Martini 3 (38,39), as the latter has been shown to underestimate the interaction strength between FGF2 and PI(4,5)P₂ (39). After 10 μs of simulation, the FGF2 hexamer induced a pronounced membrane deformation, characterized by the generation of negative curvature beneath the protein (**Figure 4 B**). Strikingly, we observed about a 4-fold enrichment of PI(4,5)P₂ molecules directly beneath the hexamer, forming a distinct nanocluster (**Figure 4 C**). Lipid enrichment and depletion around FGF2 were quantified within 1.1 nm of the protein surface using the enrichment/depletion index (40). As shown in **Figure 4 D**, palmitoyl–arachidonoyl PI(4,5)P₂ (18:0/20:4 PI(4,5)P₂; PAP2) displayed the strongest enrichment in the plasma membrane-like system (left column), leading to pronounced depletion of nearly all other lipid species, including the anionic palmitoyl– arachidonoyl phosphatidylserine (18:0/20:4 PS; PAPS). To determine whether this phenomenon depends on PI(4,5)P₂ lipid accumulation, all PI(4,5)P₂ lipids in the bulk were replaced with PAPS or palmitoyl–arachidonoyl phosphatidylcholine (18:0/20:4 PC; PAPC), but only PAPS became enriched around FGF2. To maintain biological relevance, six PI(4,5)P₂ molecules (one per monomer) were retained at experimentally validated FGF2 binding sites, ensuring stable protein– membrane anchoring. Interestingly, the polyunsaturated phosphatidylethanolamine 18:0/20:4 (PAPE) and 18:0/22:6 (PUPE) also showed moderate to strong enrichment in all systems. Both lipids contain a polyunsaturated acyl chain, which likely facilitates adaptation to highly curved membrane regions.

**Figure 4.**
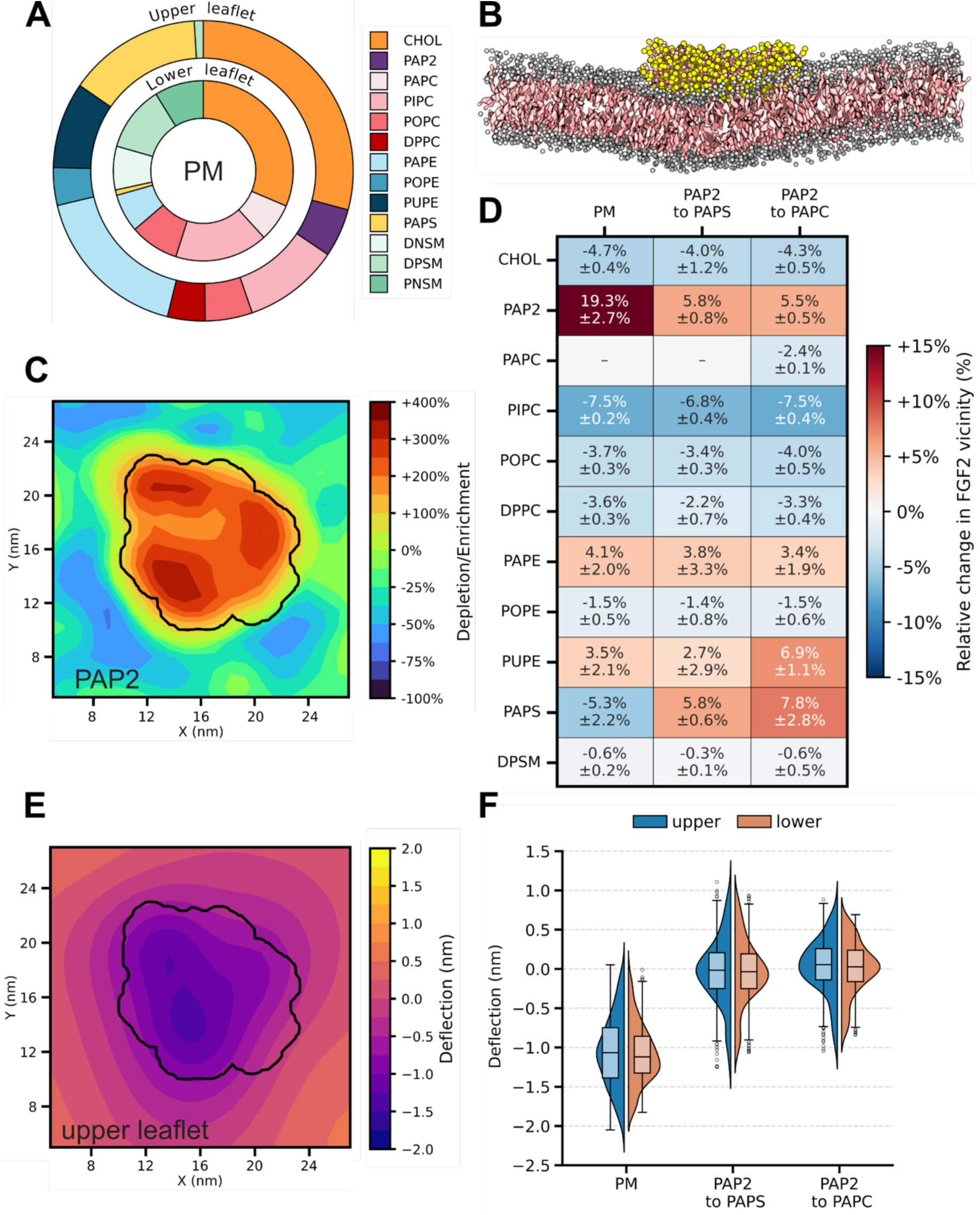
Membrane remodeling induced by PI(4,5)P₂ nanoclusters in an asymmetric plasma membrane. (A) Schematic representation of the asymmetric plasma membrane model, highlighting the lipid composition of the upper and lower leaflets. (B) Molecular dynamics snapshot showing an FGF2 hexamer (yellow) bound to the membrane surface (CHOL shown as pink surface; phosphate beads shown as gray van der Waals spheres). (C) Two-dimensional spatial deletion/enrichment map of 18:0/20:4 PI(4,5)P_2_ (PAP2) molecules in the bilayer plane relative to the center of the FGF2 hexamer. (D) Quantitative analysis (mean ± STD) of lipid enrichment and depletion in the vicinity of the FGF2 hexamer with respect to the mol% in the bulk, comparing conditions with PAP2 clusters to controls where PAP2 was replaced with PAPS or PAPC. In all cases, six PI(4,5)P_2_ molecules are bound to the FGF2 hexamer (one per monomer at the experimentally validated binding site) to preserve biological relevance. (E) Membrane deformation in the upper leaflet, shown as average bilayer height deflections induced by the FGF2 hexamer (contour lines indicate protein position). (F) Violin/box plots of membrane deflection distributions in the upper and lower leaflets under the different conditions. All analyses are based on the final 2 μs of three independent 10 μs trajectories (n = 3). Error bars represent standard deviations. Box plots indicate the median (line), interquartile range (box), and whiskers extending to the most extreme values within 1.5× the interquartile range. Outliers are shown as individual points.

Membrane deformation was further quantified by calculating leaflet height maps. The leaflet deflection was obtained by subtracting the average leaflet Z-position from the binned local Z-coordinates (**Figure 4 E**). In agreement with the snapshot shown in **Figure 4 B**, we found a pronounced downward deflection of the membrane beneath the FGF2 hexamer. Notably, this effect was absent when PI(4,5)P₂ was replaced by PS or PC, as confirmed by the deflection distributions shown in **Figure 4 F**. Together, these results demonstrate that FGF2 hexamers induce strong, PI(4,5)P₂-dependent, remodeling of the plasma membrane.

### Effect of PI(4,5)P_2_ on membrane integrity and FGF2 dependent pore formation

Based on previous mentioned data, we hypothesized that the enrichment of PI(4,5)P_2_ beneath FGF2 oligomer alters membrane properties, by weakening the bilayer, and thereby facilitating toroidal pore opening and FGF2 translocation. We evaluated the role of PI(4,5)P₂ in vesicle stability and FGF2-dependent membrane remodeling using Giant unilamellar vesicles. GUVs were prepared with POPC containing 0, 5, 10, 15, 20, and 25 mol% PI(4,5)P₂. For membrane visualization, Rhodamine-PE (Rhod-PE) was incorporated into the lipid mixture, while Alexa 647, a membrane-impermeable fluorescent dye, was added to assess membrane integrity. Membrane destabilization or pore formation was indicated by the entry of Alexa 647 into the GUV lumen (13,15,16). As shown in **Figure 5 A**, in the absence of FGF2, increasing levels of PI(4,5)P₂ led to a higher proportion of GUVs that were permeable to Alexa 647, indicating membrane destabilization at elevated PI(4,5)P₂ concentrations. At 15 mol% PI(4,5)P₂, a substantial fraction of GUVs became leaky, while above 20 mol% destabilization was even more pronounced, accompanied by reduced data reliability due to larger standard errors and fewer intact vesicles (**Figure 5 A**). These effects on vesicle yield and stability can be attributed to the intrinsic biophysical properties of PI(4,5)P₂, a non-bilayer lipid with a strong negative charge. At high concentrations, such lipids are known to destabilize bilayer structures or impair vesicle formation by introducing curvature strain and packing stress (41).

**Figure 5.**
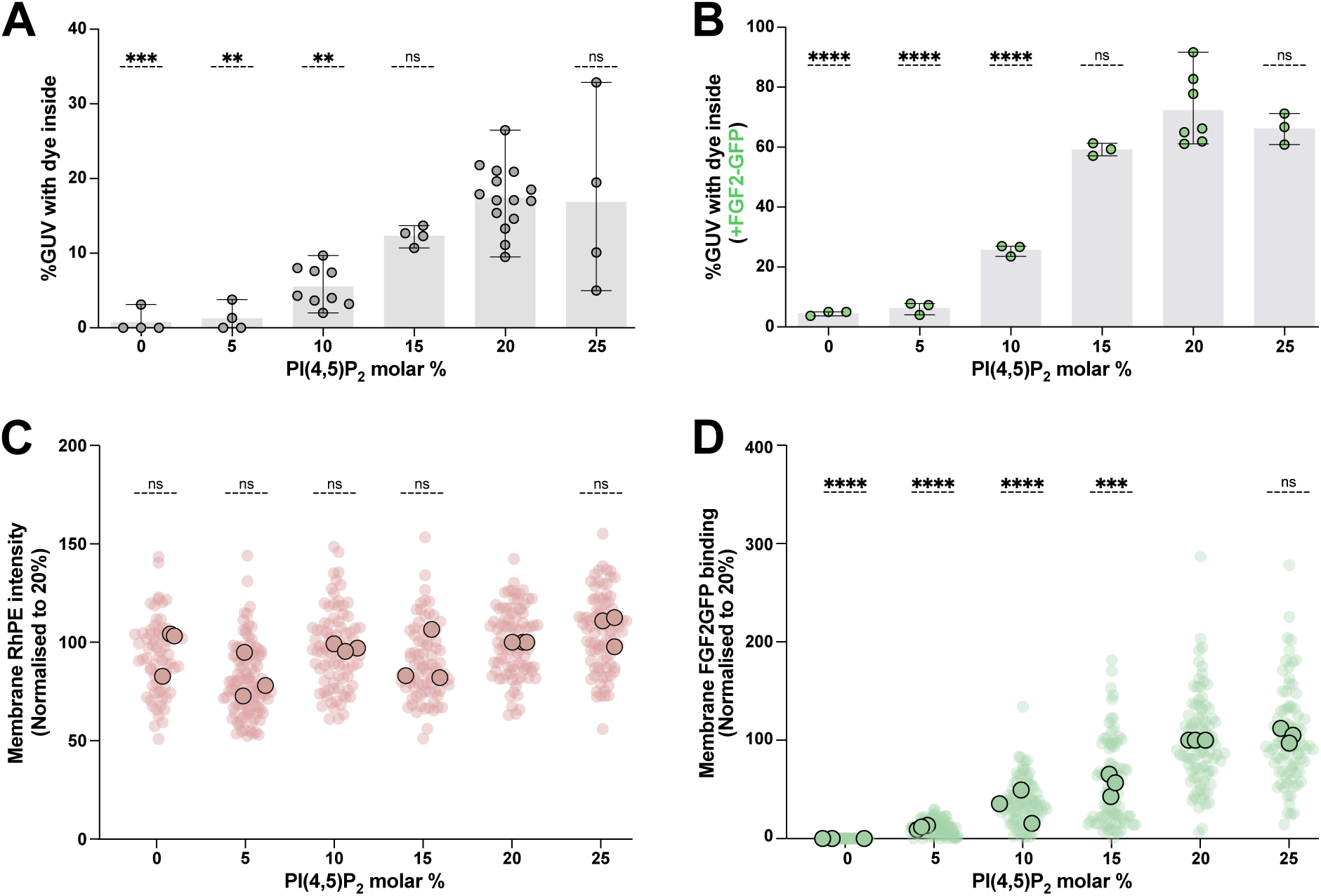
Effect of PI(4,5)P_2_ on membrane integrity and FGF2-dependent pore formation. Gaint unilamellar vesicles (GUVs) supplemented with rhodamine PE (RhPE) were prepared with increasing PI(4,5)P_2_ concentrations. (A) Quantification of GUV destabilization in the absence of FGF2 as a function of increasing PI(4,5)P₂ concentration. Each data point corresponds to a single experimental condition, in which 21-197 GUVs were analysed. Data are presented as mean with standard deviation. (B) Quantification of GUV membrane pore formation in the presence of FGF2-GFP as a function of increasing PI(4,5)P₂ concentration. Each data point corresponds to a single experimental condition, in which 20-93 GUVs were analysed. Data are presented as mean with standard deviation. (C) Membrane rhodamine PE level in GUV membranes with increasing PI(4,5)P₂ levels, measured as membrane-associated fluorescence intensity. Data is normalized to 20 mol% PI(4,5)P_2_. Each light red dot represents RhPE levels in an individual GUV from 3 replicates. Dark red dots represent mean across the three replicates. (D) Binding of FGF2–GFP to GUV membranes with increasing PI(4,5)P₂ levels, measured as membrane-associated fluorescence intensity. Data is normalized to 20 mol% PI(4,5)P_2_. Each light green dot represents FGF2-GFP binding to an individual GUV from 3 replicates. Dark green dots represent mean across the three replicates. For statistical test, data was compared with 20 mol% PI(4,5)P_2_ using ordinary one-way ANOVA in Prism. Not significant (ns) P > 0.5, **** P ≤ 0.0001. Data distribution was assumed to be normal.

To investigate FGF2-dependent membrane binding and pore formation, we employed an FGF2– GFP fusion protein. Increasing the PI(4,5)P₂ content of GUVs led to an increase in pore formation and FGF2 binding, reaching a plateau at about 15mol% to 20 mol% (**Figure 5 B and 5 D**). This indicates efficient incorporation of PI(4,5)P₂ into the bilayer up to a concentration of at least 20 mol%. As an internal quality control, we measured membrane intensity of rhodamine PE (RhPE) across different GUV populations, which remains stable (**Figure 5C**). Beyond 15 mol% threshold, the extent of FGF2-dependent pore formation remained unchanged despite a further increase in FGF2–GFP binding (**Figure 5 D**). These findings suggest that ∼15 mol% PI(4,5)P₂ represents a critical concentration at which pore formation becomes strongly favored under the given experimental conditions.

In summary, consistent with our hypothesis and MD simulations, increasing levels of the non-bilayer lipid PI(4,5)P₂ destabilized vesicles and enhanced their susceptibility to FGF2-dependent pore formation, up to a functional threshold by ∼15–20 mol%.

## Discussion

The present study builds on earlier findings indicating that C95–C95 disulfide-bridged FGF2 dimers serve as the fundamental units of higher-order FGF2 oligomers (16,42). Building on (i) brightness analyses determining the oligomeric state of FGF2 on membrane surfaces of GUVs, (ii) single-molecule TIRF microscopy quantifying the oligomeric state of FGF2 at the inner plasma membrane leaflet in living cells, (iii) extensive multiscale MD simulations to study PI(4,5)P_2_-dependent FGF2 self-assembly on membrane surfaces, and (iv) biochemical membrane integrity assays studying membrane pore formation as a function of the PI(4,5)P_2_ concentration, we propose a mechanistic model underlying membrane pore formation, the mechanistic basis of FGF2 membrane translocation into the extracellular space. Specifically, we find FGF2 oligomers to drive a local accumulation of the non-bilayer lipid PI(4,5)P_2_ which, upon passing a critical threshold concentration, causes membrane remodeling in terms of a transformation of the lipid bilayer into a toroidal membrane pore, thereby facilitating FGF2 translocation into the extracellular space (**Figure 6**).

**Figure 6.**
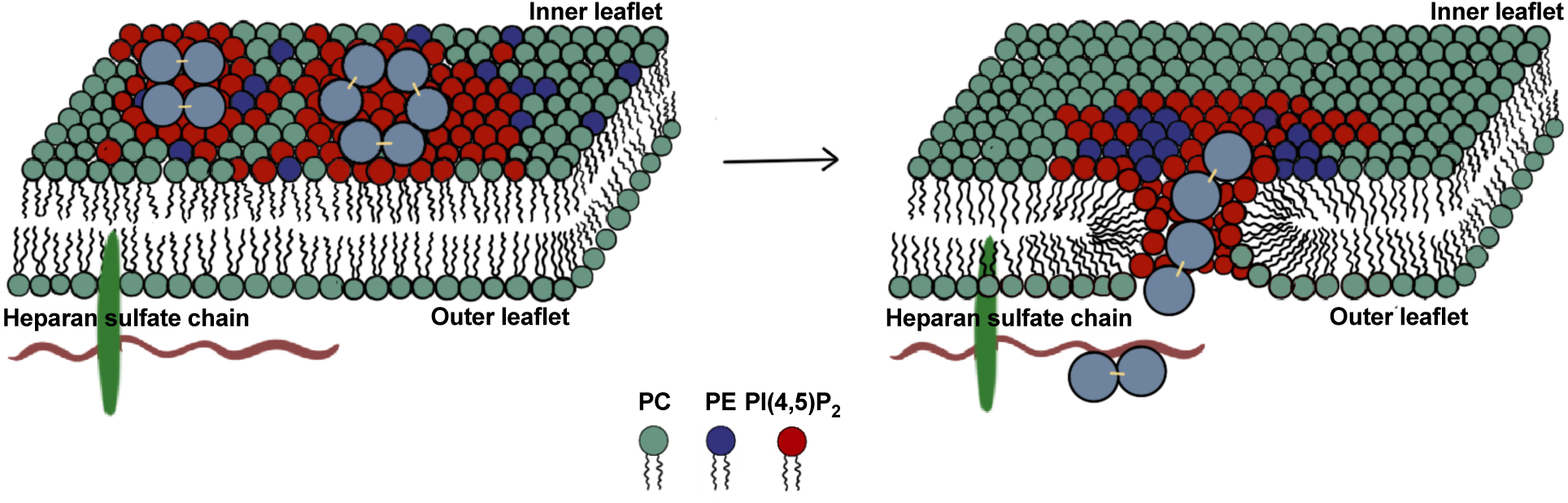
Molecular mechanism of FGF2 membrane translocation. FGF2 oligomers promote the sorting and local accumulation of non-bilayer lipids. Once this accumulation overcome a critical threshold, it triggers membrane remodeling, creating the conditions required for FGF2 translocation.

Our findings demonstrate that FGF2 predominantly assembles into tetramers and hexamers on the membrane surfaces of giant unilamellar vesicles, employing fluorescence correlation spectroscopy (**Figure 1**). Consistent with the GUV reconstitution experiments, single-molecule TIRF microscopy in living cells revealed that FGF2 predominantly assembles into tetramers and hexamers at the inner plasma membrane leaflet. Brightness analysis demonstrated that wild-type FGF2-GFP forms a dynamic range of higher-order oligomers, whereas the C95A mutant, deficient in pore formation and secretion, was restricted to dimers and monomers **(Figure 2)**. These results confirm that dimers serve as the building blocks of higher-order assemblies, but that the formation of covalent C95–C95 disulfide bridges is essential for stable oligomerization beyond the dimeric state, as previously demonstrated (15,16,42). Together with the data on GUVs, these findings establish tetramers and hexamers as the functional oligomeric species of FGF2 that mediate membrane pore formation and translocation.

Building on the experimental identification of tetramers and hexamers as the dominant functional FGF2 oligomers, extensive multiscale molecular dynamics simulations provided a structural and mechanistic framework for their self-assembly and membrane remodeling activity. Starting from C95–C95 disulfide-linked dimers, atomistic simulations revealed the spontaneous formation of ring-shaped hexamers that remained stable over extended timescales and were further stabilized by PI(4,5)P₂ molecules bound at canonical sites (K127, R128, K133) and newly formed dimer–dimer interfaces **(Figure 3)**. Coarse-grained simulations in asymmetric plasma membrane models demonstrated that such hexamers drive the local enrichment of PI(4,5)P₂ and polyunsaturated PE into nanoclusters, accompanied by bilayer deformation. Importantly, this remodeling was strictly PI(4,5)P₂-dependent, as the replacement of 18:0/20:4 PI(4,5)P₂ with 18:0/20:4 PS or 18:0/20:4 PC abolished membrane deformation. Together with biochemical reconstitution and live-cell TIRF data, these simulations establish that tetrameric and hexameric FGF2 assemblies are not only the predominant oligomeric states at membranes but also serve as structural units driving lipid sorting and membrane remodeling, as illustrated in **Figure 6**.

Finally, reconstitution experiments with defined GUV compositions established a direct role of PI(4,5)P₂ in modulating membrane integrity and FGF2-dependent pore formation. Increasing PI(4,5)P₂ concentrations progressively destabilized vesicles and enhanced FGF2 binding, with pore formation sharply increasing at a threshold of about 15 to 20 mol% PI(4,5)P₂. Beyond this level, membranes became unstable, and vesicle rupture was frequently observed.

The mechanism by which FGF2 translocate across membranes shows strong similarities to established pore-forming proteins such as Gasdermin-D. In both cases, the protein first engages with the plasma membrane through specific interactions with acidic lipids, followed by the formation of small oligomeric species (43) and subsequent oligomerization into ring-like structures (36,44–46). A key factor in this process is the local enrichment of the non-bilayer lipid PI(4,5)P₂. PI(4,5)P₂ is commonly associated with positive intrinsic curvature due to its bulky, highly charged headgroup. However, its influence on membrane bending is highly context-dependent and modulated by molecular interactions. Notably, lipidomic analyses of brain extracts reveal that the predominant PI(4,5)P₂ species contains arachidonic acid — a polyunsaturated fatty acyl chain with four double bonds. The high degree of unsaturation introduces pronounced disorder and kinking, enhancing chain flexibility and lateral mobility. These features are likely to influence the effective molecular geometry of PI(4,5)P₂ (25,47–49). These structural characteristics are believed to favor negative curvature, particularly when PI(4,5)P₂ interacts with FGF2, and we propose that this property is critical for driving FGF2 pore formation.

The non-lamellar nature of PI(4,5)P₂, combined with the closed architecture of oligomeric FGF2 (and related pore-forming proteins), appears to promote the segregation of a specialized membrane domain with distinct biophysical properties from the surrounding bilayer. When PI(4,5)P₂ reaches a critical local concentration, it alters membrane organization in the vicinity of FGF2 oligomers, inducing curvature and membrane defects. These changes lower the energetic barrier for pore opening and thereby facilitate protein translocation. Strikingly, the molecular dynamics simulations presented in this study further revealed that phosphatidylethanolamine, another non-bilayer lipid (50–52), is enriched around FGF2. This is particularly interesting given that PE constitutes about 40 mol% of the inner leaflet, which may contribute to the greater flexibility and curvature propensity of the inner plasma membrane compared to the stiffer outer leaflet (53–55). Specifically, we observed a selective enrichment of polyunsaturated PE species in the immediate vicinity of FGF2. The intrinsic negative curvature of PE, combined with the enhanced flexibility of polyunsaturated acyl chains, makes these lipids particularly well-suited to accommodate the local membrane deformation induced by protein oligomerization. Notably, polyunsaturated lipids have been shown to lower the free energy barrier for pore formation, providing a mechanistic explanation for their role in facilitating FGF2 translocation (56,57).

The combined findings from this study suggest a unifying principle by which local lipid composition and geometry critically influence pore formation. In the case of FGF2, this involves PI(4,5)P₂-driven remodeling of the inner plasma membrane leaflet. A similar mechanism operates in Gasdermin-D, which also targets PI(4,5)P₂ in the inner leaflet to nucleate oligomeric pore structures. This concept can be further generalized to other pore-forming proteins such as BAX, which binds the acidic non-bilayer lipid cardiolipin and oligomerizes into ring-like assemblies (58,59). Notably, PE is also the second most abundant phospholipid in the mitochondrial membrane (52,60), and lipid unsaturation has been shown to promote BAX and BAK pore activity during apoptosis (56), mechanisms that share striking similarities with FGF2 translocation. Beyond pore formation, the same lipid-driven principles extend to other membrane remodeling processes. For example, the SNARE complex recruits PI(4,5)P₂ and locally bend membranes (61–63), suggesting that enrichment of non-bilayer lipids may serve as a common trigger for diverse remodeling processes. Thus, we propose that the enrichment of non-bilayer lipids could represent a unifying principle that facilitates pore formation, protein translocation, and a broad range of membrane remodeling events.

## Acknowledgments

This work is supported by grants of the Deutsche Forschungsgemeinschaft (DFG) to FL, WN and KC (SFB-1638/1 – 511488495 - Z01, SFB/TRR 186, project A1, DFG LO 2821/1-1, DFG Ni 423/10-1, SFB-1557, project P1). RŠ acknowledges GAČR grant 25-15346S. PR and MH acknowledge GAČR grant 25-16481S. The authors gratefully acknowledge the data storage service SDS@hd supported by the Ministry of Science, Research and the Arts Baden-Württemberg (MWK) and the German Research Foundation (DFG) through grant INST 35/1503-1 FUGG. FL acknowledged the computing resources provided by the CSC – IT Center for Science Ltd. (Espoo, Finland) and supported by the state of Baden-Württemberg through bwHPC and the German Research Foundation (DFG) through grant INST 35/1597-1 FUGG

## Material and Methods

### Dual(+1)-FCS: Measurement

#### Preparation of giant unilamellar vesicles (GUVs) for dual(+1)-FCS: Measurement

The plasma-membrane mimicking GUVs composed of 33 mol% phosphatidylcholine (PC), 9 mol% phosphatidylethanolamine (PE), 5 mol% phosphatidylserine (PS), 5 mol% phosphatidylinositol (PI), 15 mol% sphingomyelin (SM), 30 mol% cholesterol (Chol), 1 mol% Biotinyl-PE (Avanti Polar Lipids), 0.002 mol% dioleolyl-PE labelled in the headgroup by Atto-633 (Atto-633-DOPE, ATTO-TEC) and 2 mol% phosphatidylinositol-4,5-bisphosphate (PIP2) (Avanti Polar Lipids) were prepared by electro-swelling method using platinum electrodes (64). The resulting suspension was dried under a nitrogen stream and the residual organic solvent was removed by a vacuum desiccator. The dried lipid film was resuspended with a 300 mM sucrose solution (300 mOsmol/kg). Electro-swelling was conducted at 45°C (10 Hz, 1.5 V for 50 min, 2 Hz, 1.5 V for 25 min). The GUVs were collected by centrifugation (1200x g; 25°C; 5 min) and gently washed with HEPES buffer (25 mM HEPES pH7.4, 150 mM NaCl, 310 mOsmol/kg). The washing step was repeated and the GUVs containing pellet was resuspended in a small volume of HEPES buffer. The microscopy experiments were performed in imaging chambers (IBIDI), whose surface was sequentially treated with 0.1 mg/ml Biotin-BSA (Sigma) and 0.1 mg/ml Neutravidin (Thermo Fisher Scientific) dissolved in MiliQ water. All measurements were performed at room temperature.

#### Determination of FGF2 oligomer size by dual(+1)-FCS: Measurement

The oligomer size of His-FGF2-eGFP was determined by dual(+1)-FCS (30) on a home built confocal microscope system consisting of an Olympus IX71 inverted microscope body (Olympus, Hamburg, Germany) with a 3D piezo positioner from Physik Instrumente (P-562.3CD stage controlled via E-710.3CD controller) and pulsed diode laser heads PicoTA-532, LDH-P-C-470 and LDH-P-635 controlled via PDL 828 Sepia II laser driver (all devices from PicoQuant, Berlin, Germany). The lasers were pulsing alternately to avoid artifacts caused by signal bleed-through. The beam was coupled to a single polarization maintaining single mode fibre, collimated by an air space objective (UPlanSApo 4x, NA 0.16) and directed towards the water immersion objective (UPlanSApo 60x w, NA 1.2) by a Chroma ZT375/473/532/635rpc quad-band dichroic mirror. The collected signal, which passed through a 50 micrometer diameter hole in the focal plane, was split between two SPAD detectors ($PD-50-CTC, MicroPhotonDevices, Bolzano, Italy) using T635lpxr splitter and HQ515/50 (His-FGF2-eGFP) and HQ697/58 (DOPE-Atto 633) filters mounted in front of each detector. Time-tagged time-resolved single photon counting data acquisition was performed by HydraHarp400 Multichannel Picosecond Event Timer & TCSPC Module controlled via SymPhoTime software (both from PicoQuant, Berlin, Germany). The laser intensity at the back aperture of the objective was kept below 10 mW for each laser line.

The experiment was initiated by adding His-FGF2-eGFP to a microscopy chamber containing 350 µL of GUV solution, resulting in a final His-FGF2-eGFP concentration of 200 nM. Prior to each dual(+1)-FCS measurement, individual GUVs were classified as either permeabilized or intact using Alexa 532 dye, which was added to the external solution and excited with a 532 nm laser. GUVs that were permeable exhibited Alexa 532 fluorescence in their interior, whereas intact vesicles appeared dark inside. To determine the biologically relevant oligomer size of FGF2, only permeabilized GUVs—containing membrane-inserted FGF2—were selected for FCS measurements, which were conducted within the first hour of incubation. FCS was performed at the top of a selected GUV, with the membrane positioned in the focal volume of both 470 nm and 635 nm laser beams. A 60-second dual-color FCS recording was then acquired: His-FGF2-eGFP fluorescence was detected in the first channel to assess its oligomerization, while the diffusion of the lipid marker DOPE-Atto 633 was monitored in the second channel.

#### Determination of FGF2 oligomer size by dual(+1)-FCS: Analysis

The oligomer size of His-FGF2-eGFP was determined by comparing its molecular brightness to that of a monomeric reference, Y81pCMF-C77/95A-eGFP. Dual-color FCS autocorrelation curves, *G*, were fitted using a model for two-dimensional diffusion with an additional triplet state component (65):

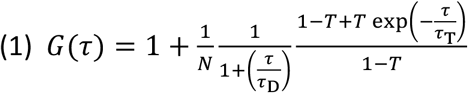

Here represents the lag-time, *N* the number of membrane-bound dye molecules in the confocal volume, is the diffusion time of membrane bound dye, SP the structure parameter, *T* the fraction of the dye in the triplet state and is the lifetime of the triplet state. Autocorrelation curves were recorded for both DOPE-Atto633 (used to confirm membrane integrity) and His-FGF2-eGFP. The FGF2 curves were analyzed to extract the molecular brightness, defined as where is the average fluorescence intensity, and *N* is obtained from the fit of the autocorrelation function. The monomeric brightness was determined using the monomeric control Y81pCMF-C77/95A-eGFP, which binds the membrane as a single unit, and by diluting His-FGF2-eGFP with nonfluorescent wild-type His-FGF2 at a 1:10 ratio to ensure that each FGF2 cluster contains at most one fluorescent emitter (15,42,66). The oligomer size of FGF2 was then calculated as the ratio of oligomer to monomer brightness:

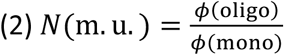

This yielded the average number of monomeric units (m.u.) per FGF2 cluster on the membrane.

### Single-Molecule TIRF Microscopy and Brightness Analysis

#### Sample preparation

CHO-K1 cells stably expressing either TMD-mGFP, FGF2-WT-mGFP or FGF2-C95A-mGFP (all constructs used in this study contained a GFP variant termed mGFP that on its own is restricted strictly to a monomeric state) in a doxycycline-dependent manner were maintained in AlphaMEM Eagle supplemented with 2 mM glutamine (PAN Biotech - P04-21250) plus 10 % FBS (PAN Biotech - P30-3031) at 37% and 5% CO_2_. Cells were detached by Trypsin/EDTA (Pan Biotech, P10-024100) treatment and seeded on glass microscopy chips. Cells were washed 3 times with PBS and fixed in 4% PFA for 20 min followed by 3 additional washing steps in PBS. Chips were maintained in PBS, stored at 4 °C and used for imaging within 24 hours.

#### Single-molecule total internal reflection fluorescence (TIRF) microscopy

Cells with low levels of FGF2-GFP expression were selected to maintain a single-molecule imaging regime and properly focused using a region of interest (ROI) of 256 x 256 pixels. Imaging was subsequently conducted in a region adjacent to the one used for selection, ensuring it had not been previously exposed to avoid measurements being compromised by bleaching. All samples were imaged on a custom-designed TIRF microscope (67) for a total of 1500 frames under a 50 ms exposure time. Excitation from a 488 nm laser at about 200 W/cm^2^ (Sapphire, Coherent) was coupled into a single mode polarization maintaining fiber to a TIRF module connected to an Olympus IX83 inverted microscope with a hardware autofocus system (IX3-ZDC, Olympus) and a 100x oil-immersion objective (UPLAPO100xOHR). Images were additionally magnified by 1.6x (IX3-CAS, Olympus) to obtain a final magnification of 160x and a pixel size of 100 nm. Fluorescence was filtered by a four-line polychroic mirror (zt405/488/561/640rpc, Chroma, 3 mm) and rejection band filter (zet405/488/561/647 TIRF, Chroma), and the emission was focused on an iXon Ultra EMCCD Camera (Andor Technologies).

#### Stoichiometry analysis

Prior to each set of experiments, we calibrated the fluorescence signal of the TIRF microscope to prevent artefacts resulting from minor adjustments in the optical setup. The calibration dataset necessary to calculate the intensity of monomeric GFP was obtained by sampling TMD-mGFP expressing cells. Images obtained from TIRF microscopy were analysed through a single-molecule brightness analysis utilizing the Stoichiometry Analysis Software (SAS) (68). Briefly, bright spots were automatically detected using an implementation of the Difference of Gaussians method combined with thresholding. The selected particles were defined using a ROI with a defined pixel size and were fitted to two-dimensional (2D) Gaussian models. The background was subtracted by defining a second ROI around the particle’s ROI with a larger pixel size. The brightness value for each detected spot was then provided by the algorithm. To avoid overlapping ROIs or multiple particles in the same ROI, localized particles were filtered based on the distance between two ROIs. The distribution of all the brightness values was plotted as a kernel probability density function (Pdf). The mean monomer intensity and standard deviation, calculated in the calibration phase from monomers of the GFP dataset, were used to define the theoretical intensity of possible higher-order oligomeric species. This approach generated a linear combination of Gaussians, which was then used to fit the experimental Pdf. Percentages of occurrence of oligomeric species of FGF2 were calculated from the area of each fitted Gaussian. Values were corrected for partial fluorescence efficiency of GFP. Detection parameters were configured as follows in SAS: camera pixel size (nm): 100, camera quantum efficiency: 95, camera offset: 170, camera EM gain (counts per photoelectron): 65.4, maximum sigma: 200. Analysis parameters were set as: labeling efficiency (for GFP): 70, maximum Gaussian mixtures: 30 (max).

### Multiscale Molecular Dynamics Simulations

#### Force Fields and Simulation Software

All molecular dynamics simulations were carried out using GROMACS 2023 for atomistic systems and GROMACS 2024 for coarse-grained systems. (35). Atomistic simulations employed the CHARMM36m force field (February 2021 release) for proteins, the CHARMM TIP3P model for water, and the CHARMM36 parameters for ions (34,69). For coarse-grained simulations, the Martini 2.2 (37) force field was used, applying an optimized bond-constraint topology for cholesterol (70).

##### Protein System Preparation

The dimeric structure of FGF2 was derived from previous modeling, integrating molecular dynamics simulations, cryo-EM density, mass-spectrometry cross-linking data, and AlphaFold predictions (16). Shortly, this structure was built from the monomeric crystal structure (PDB ID: 1BFF (71)) covering residues 26–154 of FGF2. Two monomers were connected via a C95–C95 disulfide bond with neutral termini (–NH₂, –COOH) to prevent non-specific electrostatic interactions that could hinder sampling. Two 18:0/20:4 PI(4,5)P₂ molecules were pre-docked (taken from existing simulations (16)) into the experimentally validated binding pocket residues (K127, R128, K133). The resulting dimer was inserted into pre-built membranes using a lateral pressure insertion method following M. Javanainen protocol (72). Atomistic membranes were generated with the CHARMM-GUI Membrane Builder (73), while CG membranes were prepared using TS2CG 2.0 (74,75).

#### Atomistic Molecular Dynamics Simulations

For self-assembly studies, 27 independent systems were constructed, each containing three dimers placed in distinct orientations with a minimum inter-dimer separation of 1 nm to allow weak intermolecular interactions (van der Waals and Coulomb cutoffs = 1.2 nm. Simulations were run for at least 3.9 μs each, yielding a cumulative sampling of 138.25 μs (see **Table 2** for details). An additional 1.6 μs simulation was performed with a third dimer positioned near a preformed V-shaped tetramer. To assess the stability of the ring-like hexamer, the oligomer was embedded in a complex asymmetric plasma membrane composition adapted from Schaefer and Hummer (36), but with 18:0/20:4 PI(4,5)P₂ lipids matching experimental conditions and lower cholesterol concentration (30 mol%) to reduce membrane rigidity and enhance lipid diffusion (see **Table 3** for details). For this setup, three independent simulations of 5 μs each were conducted with different initial velocity seeds to ensure statistical independence. Protein–membrane systems were energy-minimized in vacuum, solvated with TIP3P water, neutralized with counter-ions, and supplemented with 150 mM KCl to match experimental salt concentration. Energy minimization was followed by equilibration in the NpT ensemble with position restraints applied to all protein atoms and the first heavy atom of each lipid (the protein was position-restrained with a force constant of 1000 kJ/mol, applied in all directions, while the lipids were restrained only in the xy-plane). Temperature was controlled at 310 K using the Nosé–Hoover thermostat (76) (τ = 1.0 ps), and pressure was maintained at 1 bar with the Parrinello–Rahman barostat (77) (τ = 5.0 ps; compressibility = 4.5 × 10⁻⁵ bar⁻¹). Semi-isotropic pressure coupling was applied. Electrostatic interactions were computed with the Particle Mesh Ewald method (78), and neighbor searching employed the Verlet cutoff scheme (79) with updates every 20 steps. Simulations used a 2 fs timestep with periodic boundary conditions in all directions.

#### Coarse-Grained Molecular Dynamics Simulations

For CG systems, protein dimers were converted using martinize with vermouth v0.9.6 (80), applying an elastic network to each dimer (force constants: –ef 700.0; lower/upper distance bounds: –el 0.5, –eu 0.9). We used two different membrane types. The the asymmetric plasma membrane–like composition where the composition was adapted from the work of Schaefer and Hummer (36) but adjusted due to Martini 2 lipidome limitations (see **Table 4** for details). Three different membrane compositions were employed. The first was an asymmetric plasma membrane–like system, adapted from S. Schaefer and G. Hummer (36) and adjusted to account for limitations of the MARTINI 2 lipidome (see **Table 4** for details). The others are a modified system in which PI(4,5)P_2_ (PAP2) lipids were converted to PAPS to PAPC, excluding PAP2 molecules pre-bound to FGF2. Each system was simulated in triplicate for 5 μs, and unless otherwise noted, analyses were performed on the final replicate. Independent trajectories were generated by randomizing initial velocities. Electrostatics were treated with the reaction-field method (rcoulomb = 1.1 nm, εr = 15, εrf = 0). Van der Waals interactions used a cutoff scheme with potential-shift-Verlet modification and rvdw = 1.1 nm. Temperature was maintained at 310 K using the v-rescale thermostat (81) (τ = 1.0 ps, coupling every 20 steps). Pressure was controlled semi-isotropically with the Parrinello–Rahman barostat (τ = 12.0 ps, compressibility = 3 × 10⁻⁴ bar⁻¹, reference pressure = 1 bar). To reduce neighbor list artifacts, we set verlet-buffer-tolerance = -1 and rlist = 1.35 nm (82).

**Table 4.** Initial lipid compositions of the asymmetric plasma membrane–like system (PM), represented using MARTINI 2.2 lipid nomenclature. Values are adapted from Schaefer and Hummer (36). Three membrane setups are shown: (i) the reference PM system, and (ii) a modified system in which PI(4,5)P_2_ (PAP2) lipids were converted to PAPS or (iii) PAPC, excluding PAP2 molecules pre-bound to FGF2. Approximately 100 cholesterol molecules located in the membrane midplane were evenly redistributed between the two leaflets.

| Acyl chain and type | Martini 2 names | PM upper | PM lower | PAP2 to PAPS upper | PAP2 to PAPS lower | PAP2 to PAPC upper |
| --- | --- | --- | --- | --- | --- | --- |
| Cholesterol | **CHOL** | 454 | 516 | 433 | 537 | 433 |
| 18:0 - 20:4<br>PI(4,5)P<sub>2</sub> | **PAP2** | 81 | 0 | 6 | 0 | 6 |
| 16:0 - 20:4-PC | **PAPC** | 0 | 113 | 0 | 113 | 75 |
| 18:2 - 16:0-PC | **PIPC** | 160 | 275 | 160 | 275 | 160 |
| 18:1 - 18:0-PC | **POPC** | 79 | 145 | 79 | 145 | 79 |
| 16:0 - 16:0-PC | **DPPC** | 63 | 0 | 63 | 0 | 63 |
| 16:0 - 20:4-PE | **PAPE** | 271 | 113 | 271 | 113 | 271 |
| 18:1 - 18:0-PE | **POPE** | 63 | 0 | 63 | 0 | 63 |
| 18:0 - 22:6-PE | **PUPE** | 144 | 0 | 144 | 0 | 144 |
| 18:0 - 20:4-PS | **PAPS** | 224 | 16 | 299 | 16 | 224 |
| d18:1 - 24:1-SM | **DNSM** | 0 | 129 | 0 | 129 | 0 |
| d18:1 - 18:0-SM | **DPSM** | 15 | 194 | 15 | 194 | 15 |
| d18:1 - 24:0-SM | **PNSM** | 0 | 145 | 0 | 145 | 0 |

**Table 5.**
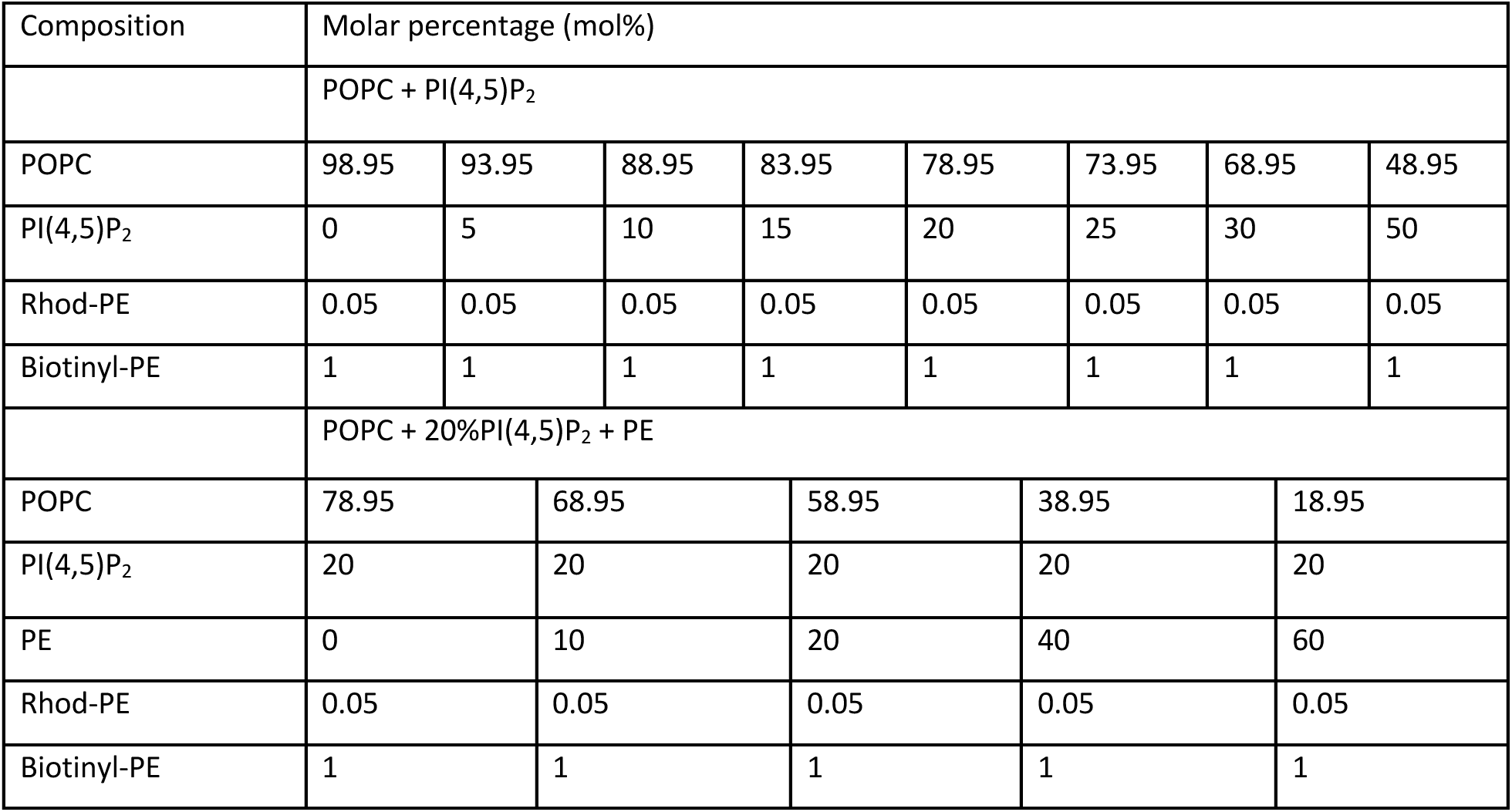
Lipid compositions of GUVs used for vesicle stability and pore formation assay.

#### Analysis

All analyses were conducted using an in-house tool in combination with MDAnalysis (83) and lipyphilic (84).

##### Lipid depletion/enrichment around FGF2 Hexamer

To quantify the local enrichment or depletion of lipid species around a protein within a cutoff of 1.1 nm, we compute the depletion/enrichment index (DEI) as:

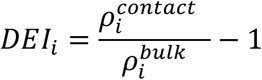

where 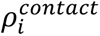 is the relative abundance (or normalized number density) of lipid species *i* in contact with the protein, and 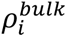 is the corresponding abundance of lipid species *i* in the bulk membrane (outside of the contact cut-off).

##### Surface deflection

The leaflet deflection is obtained by subtracting the average Z position of the leaflet from the binned Z positions, as determined by the positions of the lipid phosphate beads, giving an estimate of the membrane deformation.

#### Vesicle Stability and Pore Formation Assay

##### GUV Preparation

All lipids were purchased from Avanti Polar Lipids (850457P-POPC, 840046P-PI(4,5)P_2_, 840026C-PE, 810158P-RhodPE, 870282C-BiotinylPE) and stored in chloroform at -20°C under Argon. Protocol for GUV generation via electroformation method was adapted from (15) with minor modifications. Lipids were mixed in chloroform at above indicated compositions and the resulting lipid mixture was stored and used within 1 week after its preparation. Volume of 5μl of the lipid mixture were spread on two platinum electrodes and dried for 10-15mins. The electrodes were then immersed in chambers containing 350μl of 300mM sucrose solution (osmolality: 301mOsmol/kg). Osmolality was determined with Wescor Vapro 5600 instrument. GUV formation was induced by swelling at 45°C for 50mins (10Hz, 1.5V) followed by 25mins at 2Hz (1.5V) to facilitate detachment of GUVs from the electrodes. After gradually cooling the GUVs to room temperature for 20mins, they were gently washed with 150mM KCl, 25mM HEPES-KOH, pH 7.34 (osmolality: 308mOsmol/kg) buffer (HK buffer) via centrifugation at 1200xg, 25°C for 5mins. The supernatant was removed to leave ∼500μl buffer in which the loose GUV pellet was resuspended. For imaging we used, µ-Slide 8 Well^high^ Glass Bottom chambers from Ibidi (80807). To immobilize the GUVs on the glass surface, imaging chambers were blocked with 0.1mg/ml Albumin, biotin labeled bovine (Biotin-BSA, A8549, Sigma Aldrich) followed by 0.1mg/ml Avidin, NeutrAvidin™ Biotin-binding Protein (Neutravidin, A2666, Invitrogen^TM^). In between the blockings, the chambers were washed with water and at the final step, with HK buffer. 100µL GUVs and 100µL HK buffer was added in the ibidi chambers. For testing vesicle stability, GUVs were incubated with small fluorescent tracer-Alexa647 (A33084, Invitrogen) for 20mins. For FGF2 binding assay, the GUVs were incubated with Alexa647 and 200nM FGF2-GFP fusion protein for 60mins.

Imaging was carried out with Zeiss LSM800 confocal microscope using 63 x/1.4 oil immersion objective lens. Images were acquired in the multitrack mode at room temperature. The samples were excited with an argon laser (488nm), and He-Ne lasers (561nm and 633nm) for GFP, Rhodamine-PE and Alexa647 respectively. Images were taken in 8-bit as a snap or tile scan mode.

##### GUV Quantification for leakage of alexa647 and FGF2GFP binding

For quantification, ImageJ software (http://rsbweb.nih.gov/ij/) was used as indicated in Steringer *et al.*, 2017. A small circle was drawn with the circle selection tool in the lumen at the center of the GUV and the immediate surrounding of the GUVs. To assess membrane pore formation, a threshold was established based on the ratio of internal to external Alexa647 tracer fluorescence signal intensity. In case of the ratio being 0.6, the vesicle was considered to have either dye inside due to membrane damage (read out of membrane integrity) or membrane pore. In order to quantify FGF2 binding, the plugin ‘radial profile angle’ (http://rsbweb.nih.gov/ij/plugins/radialprofile.html) was utilized. A big circle was drawn with the circle selection tool around the GUV. With the radial profile angle plugin along the radius of the circle, the intensity is measured at all points and processed into a profile plot of normalized integrated intensities around homocentric circles as a function of distance from the center. For each experimental condition, 20-197 individual GUVs were analyzed.

